# Early life trauma leads to escalated aggressive behavior and its inheritance by impairing thyroid hormone availability in brain

**DOI:** 10.1101/2021.07.05.448713

**Authors:** Rohit Singh Rawat, Aksheev Bhambri, Muneesh Pal, Avishek Roy, Suman Jain, Beena Pillai, Arpita Konar

## Abstract

Escalated and inappropriate levels of aggressive behavior referred to as pathological in psychiatry can lead to violent outcomes with detrimental impact on health and society. Early life trauma triggers adulthood violence and criminality, though molecular mechanisms remain elusive. Here, we provide prefrontal cortex and hypothalamus specific transcriptome profiles of peripubertal stress (PPS) exposed Balb/c adult male mice exhibiting escalated aggression and adult female mice resilient to such aberrant behavioral responses. We identify transthyretin (TTR) as a key regulator of PPS induced escalated aggression and its intergenerational inheritance. TTR mediated long-term perturbation in hypothalamic thyroid hormone (TH) availability contributed to male aggressive behavior without affecting circulating hormone. Ttr gene ablation in hypothalamus impaired local TH signaling including levels of TH transporters (Mct8, Oatp1c1), deiodinase 2(DIO2) and TH responsive genes (Nrgn, Trh and Hr). Escalated aggressive behavior and impaired TTR-TH signaling was also inherited in F1 male progenies. Further, we deciphered Ttr promoter hyper methylation in hypothalamus of such abnormally aggressive males across generations. Interestingly prefrontal cortex showed opposite pattern of Ttr expression as well as long term epigenetic changes. Also, T4 increase by levothyroxine in PFC did not produce any behavioral changes. Our findings reveal that trauma during puberty trigger lasting escalated aggression by epigenetic programming of TTR and consequent impaired thyroid availability in brain. TTR-TH signaling in brain can serve as potential target in reversal of escalated aggression and related psychopathologies.

## Introduction

Escalated aggressive behavior transforming into violent outbursts is a complex personality trait that intersects with several psychopathologies and can lead to antisocial and criminal activities (*1*). Every year, more than 1 million people worldwide die because of assault and many more are victimized of domestic violence and other forms of physical injuries. Besides afflicting the common mass, violence poses enormous financial burden for emerging society and is a major challenge to human welfare. Such global threat to humanity necessitates identification of predisposing factors and early intervention strategies. Escalated aggressive behavior marked by an inability to conform to social norm is considered pathological and indicative of violence. Normal aggression is a behavioral response to threat and competition, but when expressed out of proportion, control and context including misjudging age, sex of the opponent loses its social communicative nature. Such uncontrolled excessive aggression devoid of inhibitory mechanisms can have injurious consequences and referred to as pathological (*2*, *3*). Animal aggression can also be escalated with pathological signs, if there is a response surpassing species-typical levels; attacks targeted on inappropriate partners, and body parts prone to serious injury; attacks not signaled by threats; or ignorance of signals of opponents (*4*) In general, these criteria resemble human aggressiveness expressed in certain psychopathologies.

The key to combat violent behavior is deciphering the triggers underlying brutal shift of normal adaptive aggression to escalated and pathological form. Mounting epidemiological evidences link early life traumatic experiences with adult aggression and criminality. A landmark study of 50 violent offenders with history of childhood abuse pioneered the concept that brain is susceptible to stress during critical periods of early life deteriorating mental health. In particular, trauma around puberty or adolescence including fear, maltreatment, physical and sexual abuse confers susceptibility to violence in adult individuals. Moreover, such behavioral anomalies are not limited parental generations and can be faithfully transmitted to progenies who have never been exposed to trauma. Although pathological aggression has emerged as a consequence of early life adversities, biological insights are obscure. Majority of research in the field of aggressive biology have focused on the adaptive form without really considering the excessive or inappropriate forms and clinical importance of targeting violent individuals (*5*). Essentially the lacunae in biologically relevant and valid animal models for pathological aggression are the primary reason for the gap in knowing the biological roots. Márquez et al. (*6*) developed a novel animal model which showed the effect of peripubertal fearful exposures on male pathological aggression at adulthood in Wistar Han rats. They primarily focused on neural circuits of aggression and on a single gene MAOA in isolation.

Considering multi-factorial etiology of escalated and pathological aggressive behavior, we rationalized that unbiased genome wide investigation would decipher key molecular pathways that can be exploited further as prediction and intervention targets. We modeled PPS induced escalated aggression in laboratory bred Balb/c mice and screened the male cohort showing extreme phenotypes. We selected the extreme phenotypes for better understanding of the behavior observed in human violent offenders and psychopathy. Female mice showed resilience towards peripubertal trauma induced escalated aggressive phenotype as also reported previously (*7*).

Next, we performed a sex specific transcriptome analysis in vulnerable brain regions of hypothalamus and prefrontal cortex (PFC). Hypothalamus is an integral brain region for expression of both normal and deviant or maladaptive form of aggressive behavior. Further, neural circuit specific manipulation experiments revealed that ventromedial hypothalamus is the key region for inter-male aggression (8,9) While hypothalamus is considered as the trigger center for aggression, PFC plays opposite regulatory role being involved in inhibition of threat provoked aggressive behavior. More importantly, direct neuronal projections from PFC to hypothalamus have been suggested to control both type and amplitude of aggressive behavior (*10*,*11*). Therefore, we primarily focused on hypothalamic molecular culprits of escalated aggression and also included PFC in our study to understand inter-brain regional molecular regulation if any.

We prioritized Ttr gene given its i) top rank in hypothalamus transcriptome analysis and unique sex specific diametrically opposite expression pattern in hypothalamus and PFC and iii) long term gene expression changes from early peripubertal age till adulthood. Next, we deciphered a molecular mechanism whereby PPS incited sustained TTR deficiency in hypothalamus resulted in altered levels of other thyroid hormone (TH) transporters (Mct8, Oatp1c1) and deiodinase (DIO2), reduced local TH availability and modulated expression of TH regulated genes (Nrgn, Trh) that eventually led to emergence of escalated aggressive behavior. These behavioral and molecular deficits were also transmitted to the non-stressed F1 male progenies of PPS F0 males showing escalated aggression. Further, epigenetic analysis revealed that methylation mark in Ttr promoter might attribute to such long term programming of behavior.

## Results

### Screening of escalated aggressive phenotypes

The goal of the experiment was to screen the adult animals exhibiting escalated aggressive phenotype with pathological signs in response to PPS exposure. We performed behavioral test in 5 independent cohort of mice and accordingly 5 independent experiments. Behavioral scoring and screening from these 5 cohorts were pooled and represented in Fig 1. Screening parameters were optimized based on earlier reports (12). We observed that in adult control Balb/c mice [Ctrl-RI; Total N=60 {N=15 (cohort 1)+N= 12 (cohort 2)+N=9 (cohort 3)+N=12 (cohort 4)+N=12 (cohort 5)}], 95% (N=57) were non-aggressive (Nagg) while 5% (N=3) showed normal offensive aggression (Lagg) but not escalated and devoid of any pathological signs. Amongst PPS adult male mice cohort (PPS-RI) of 60{N=15 (cohort 1)+N= 12 (cohort 2)+N=9 (cohort 3)+N=12 (cohort 4)+N=12 (cohort 5)}], 78.33% (N=47) showed abnormal aggression with signs of pathological forms characterized by prolonged fighting with short attack latency, attack on females and anesthetized intruder in all the sessions tested. Amongst these 47 mice, 32 (53.33%) showed extreme phenotypes with greater than 80% attack duration, very short attack latency of less than one minute in all the sessions (Fig. 1B,1C and Source File 1C) and those which attacked both females and anaesthetized intruder. These were referred to as escalated aggressive “Eagg” and rest as hyper-aggressive “Hagg” (N=15; 25%). We selected “Eagg” mice cohort for further molecular and cellular analyses. Some mice of the PPS-RI cohort were moderate-aggressive (Magg; N=8; 13.33%) showing signs of pathological form in some days of the RI session and 8.33 % (N=5) showed normal offensive aggression across 7 days of 10 min screening sessions. As reported earlier (*7*) females did not show escalated aggression.

**Fig. 1.**
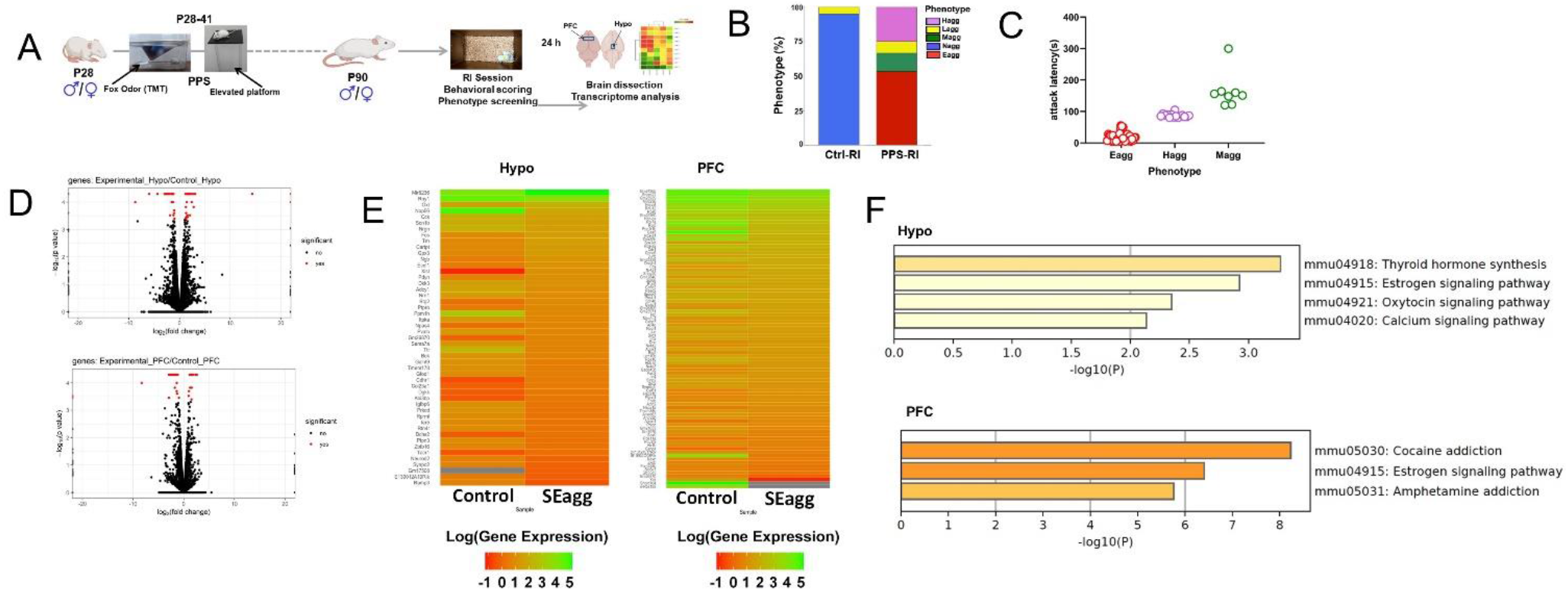
Brain region specific transcriptional responses in peripubertal stress induced adult males showing escalated aggression. (A) Experimental timeline of peripubertal stress (PPS) exposure, resident intruder (RI) behavioral paradigm, brain dissection and transcriptome analysis. (B) (i) Phenotypic behavioral screening post RI scoring in control mice without PPS exposure (Ctrl-RI; Total N=60 from 5 cohort of mice used in 5 independent experiments; N=15, N=12, N=9, N=12 and N=12) and Experimental mice with PPS exposure (PPS-RI; Total N=60 from 5 cohort of mice used in 5 independent experiments; N=15, N=12, N=9, N=12 and N=12). Histogram represents non-aggressive (Nagg; N=57) and less aggressive (Lagg;N=3) mice in the Ctrl-RI cohort. PPS-RI cohort comprises of escalated aggressive with (Eagg; N=32), hyper-aggressive (Hagg; N=15), moderate-aggressive (Magg; N=8) and less-aggressive (Lagg; N=5) mice (C) Attack latency of Eagg, Hagg and Magg mice of PPS-RI cohort. (D) Volcano plot, (E) heatmap of differentially expressed genes {DEGs; Ctrl-RI (Control) vs PPS-RI escalated aggressive (SEagg) males} in hypothalamus (Hypo) and prefrontal cortex (PFC) and (F) KEGG gene enrichment analysis in males. RNA sequencing libraries were prepared from N=3 mice per group considered as three biological replicates.

### Transcriptome analyses identify brain region specific gene signatures in PPS induced adult males showing escalated aggression and resilient females

To discover unbiased molecular correlates of early life trauma induced escalated aggression and its sex differences we used RNA-sequencing to measure all polyA-containing transcripts in hypothalamus (Hypo) and PFC of control and PPS and male and female mice. Now we refer to the peripubertally stressed adult escalated aggressive male mice group as “SEagg” and adult non-aggressive female mice group as “SNagg”. Experiment was performed in 3 individual mice considered as biological replicates for all the samples and tissues were collected 24 h after last RI session. Heatmaps of differentially expressed genes (DEG)s were constructed from the global transcriptome analysis. In Hypo, 49 genes were differentially expressed amongst which 28 were down-regulated, 20 were up-regulated and 1 was expressed in experimental males but not in control males. PFC of SEagg males showed 87 DEGs amongst which 57 were downregulated, 28 were upregulated and 2 were only expressed in experimental males but not control males (Fig. 1D, 1E)

Resilient non aggressive females (SNagg) showed more DEGs (Hypo-684;PFC-152) than SEagg males when compared to their respective control samples (Fig. 2A, 2B) Comparative analysis of male vs female showed both overlapping and discrete gene signatures. In hypothalamus, 16 DEGs overlapped between male and female, 12 showing expression changes in opposite direction, 4 in similar direction and 33 genes were exclusive in SEagg male cohort (Fig. 2D). In PFC, 15 DEGs overlapped between male and female all showing expression changes in similar direction and 72 genes were exclusive to SEagg males.

**Fig. 2.**
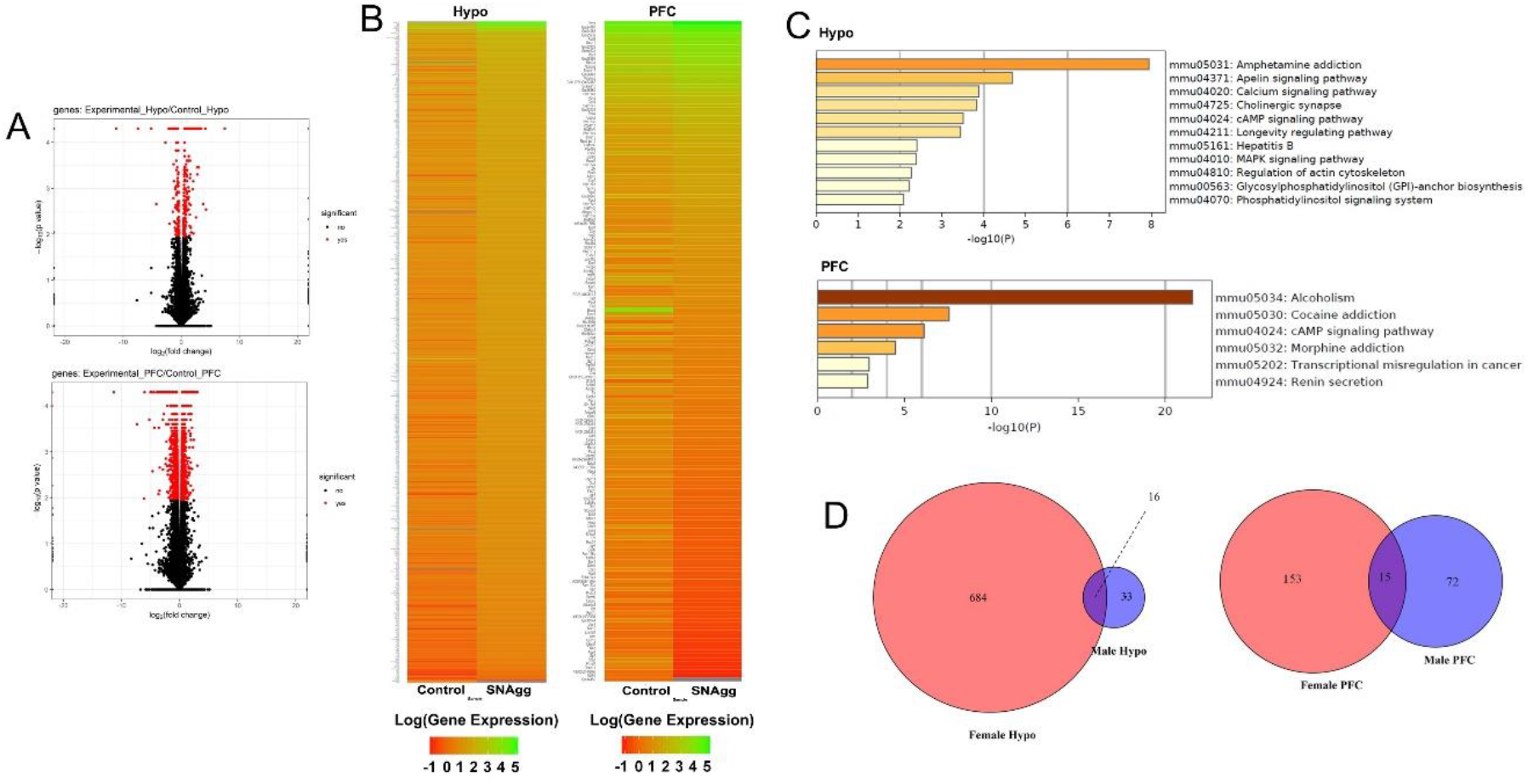
Brain region specific transcriptional responses in peripubertal stress induced adult resilient females. (A) (i) Volcano plot, (B) heatmap of differentially expressed genes (DEGs; Ctrl-RI (Control) vs PPS-RI Non aggressive (SNAgg) females} in hypothalamus (Hypo) and prefrontal cortex (PFC) and (C) KEGG analysis and (D) Venn diagram of Hypo and PFC specific overlapping DEGs between SEagg males and SNAgg females. RNA sequencing libraries were prepared from N=3 mice per group considered as three biological replicates.

In order to identify the gene signatures causal for PPS induced male escalated aggression, we prioritized genes of 2 categories including i) male exclusive DEGs in Hypo and PFC ii) DEGs that showed opposite pattern in both sexes. Amongst these DEGs, we selected top ranking 10 genes from each category and finally 20 DEGs got validated by RT-PCR.

Ttr, encoding for thyroid hormone (TH) transporter protein was the topmost ranking gene in Hypo of our transcriptome data (Fig.1D and Supplementary Fig. S1) that was validated by RT-PCR. Further, it was the only gene showing unique brain region and sex specific diametrically opposite pattern (Supplementary Fig. S1 and Fig 3). Gene ontology enrichment analysis using KEGG tool combined with literature mining also showed TH signaling as one of the top ranking pathways (1F). TH signaling genes Nrgn and Trh was amongst the top ranking genes in Hypo (Supplementary Fig S1) and showed sex specific opposite pattern in Hypo (Fig. 4).

**Fig. 3.**
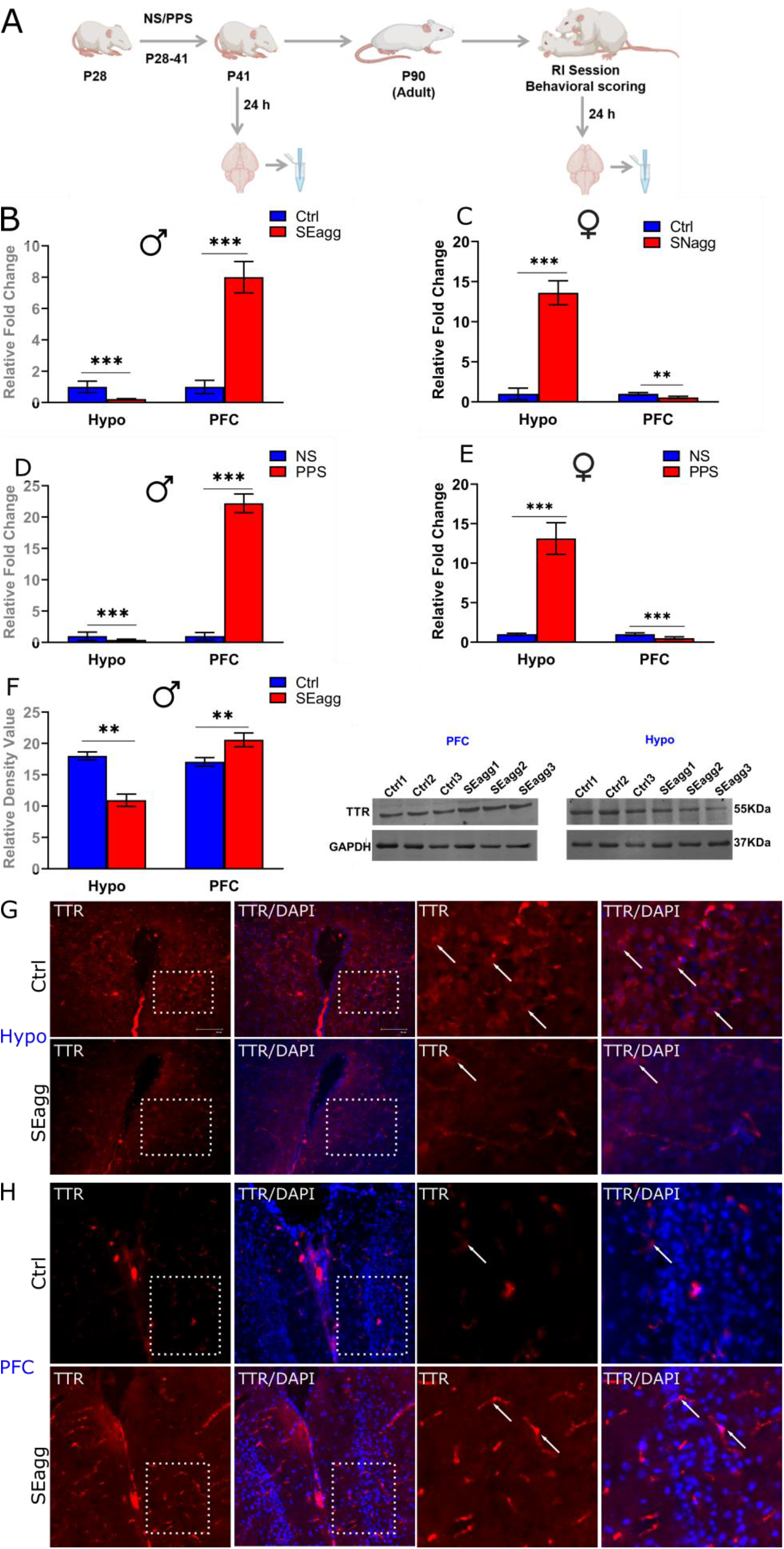
Peripubertal stress induced long term changes in TTR expression in brain region and sex specific diametrically opposed pattern. (A) Experimental timeline for transthyretin (TTR) expression analysis. Ttr mRNA expression profile in Hypo and PFC of peripubertal stress exposed (PPS) adult (B) male (SEagg) and (C) female (SNAgg) mice with control (Ctrl) counterparts 24 h after RI session (N=12 mice/group from 3 independent experiments and cohorts of mice). Ttr mRNA expression profile in Hypo and PFC of peripubertal (D) male and (E) female mice 24 h after stress exposure (PPS) with control [no stress exposure (NS)] counterparts (N=12 mice/group from 2 independent experiments and cohorts of mice). Ttr protein expression profile (F) immunoblot (N=3 mice per group and experiment repeated thrice with same samples) and immunofluorescence analysis (N=3 mice/group and experiment repeated thrice with different tissue sections of same animal) in (G) Hypo and (H) PFC of Ctrl and Eagg males. Representative Immunoblot from a single experiment with 3 mice/biological replicates each of Ctrl and SEagg groups. Histogram represents mean of the data (±SD). Statistical analysis were performed using unpaired Student’s t-test [** (p< 0.01) and *** (p< 0.001)] between NS vs PPS groups, Ctrl vs SEagg or Ctrl vs SNAgg.

**Fig.4.**
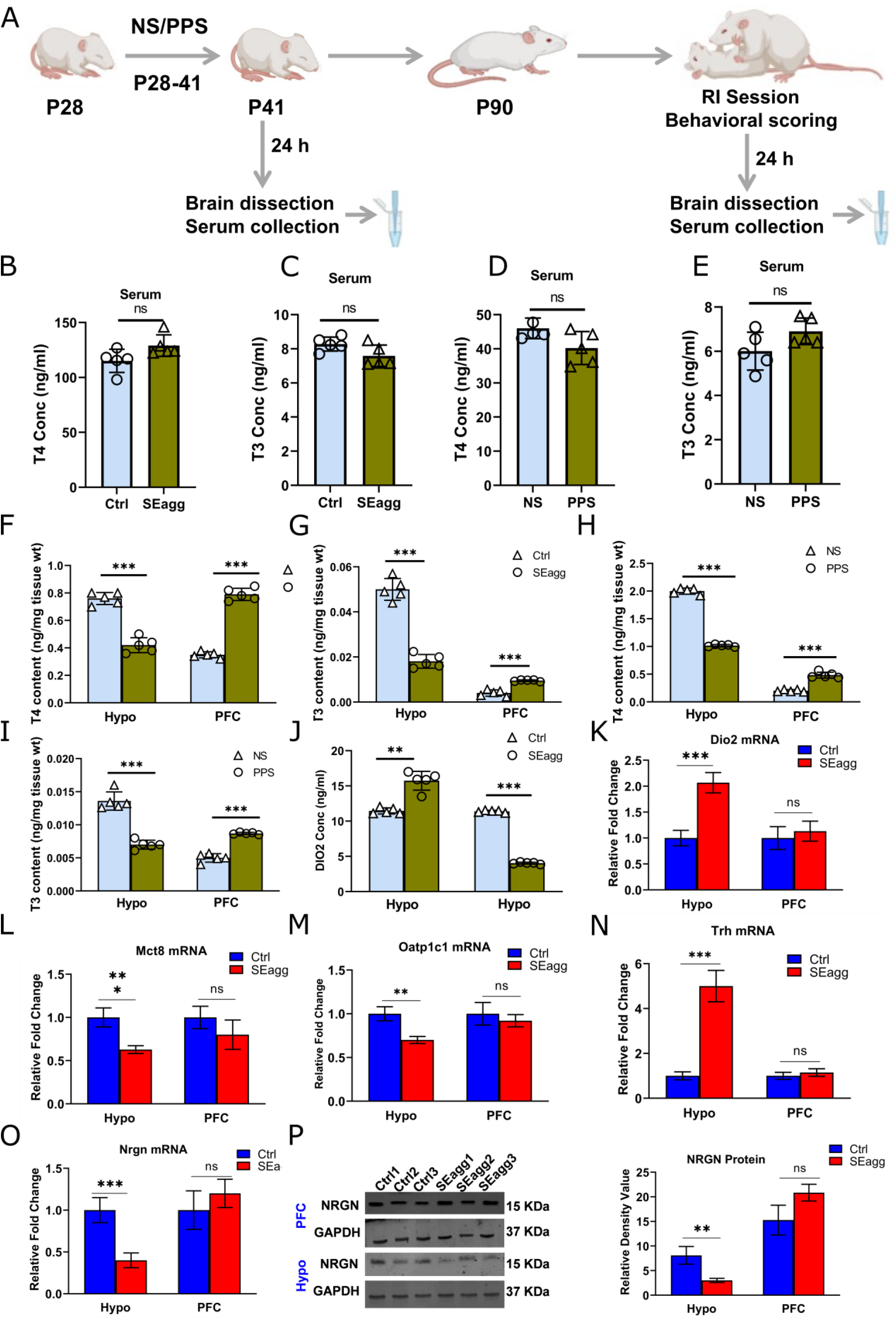
Peripubertal stress induced long term perturbation of thyroid hormone availability in the brain of escalated aggressive males with concomitant changes in transporters, deiodinase and target gene expression. (A) Experimental timeline. T4 (B) and T3 (C) level in serum of peripubertal stress exposed (SEagg) and control (Ctrl) adult males 24 h after RI session. T4 (D) and T3 (E) level in serum of peripubertal males 24 h after stress exposure (PPS) with control [no stress exposure (NS)] counterparts. T4 (F) and T3 (G) level in Hypo and PFC of peripubertal stress exposed (SEagg) and control (Ctrl) adult males 24 h after RI session. T4 (H) and T3 (I) level in Hypo and PFC of peripubertal males 24 h after stress exposure (PPS) with control [no stress exposure (NS)] counterparts. DIO2 enzyme (J) concentration in Hypo and PFC of Ctrl and SEagg males. All the above ELISA assays “B-I” were performed in N=5 mice per group from 2 independent experiments and cohorts of mice. Dio2 mRNA (K), Mct8 mRNA (L), Oatp1c1 mRNA (M), Trh mRNA (N) and Nrgn mRNA (O) expression profile in Hypo and PFC of Ctrl and SEagg males (N=9 mice per group from 3 independent experiments and cohorts of mice). NRGN protein expression (P) profile in Hypo and PFC of Ctrl and SEagg males (N=3 mice per group and experiment repeated thrice with same samples). Representative Immunoblot from a single experiment with 3 mice/biological replicates each of Ctrl and SEagg groups. Histogram represents mean of the data (±SD). Statistical analysis were performed using unpaired Student’s t-test [ns (p> 0.05), ** (p< 0.01) and *** (p< 0.001)] between NS vs PPS groups or Ctrl vs SEagg.

Amongst rest of the 17 genes (Supplementary Fig. S2), 14 were male exclusive but altered either in Hypo (downregulated Nrn1, Rtn4r, NeuroD2, Zbtb16, Pvalb; up-regulated Cartpt, Gm17508, Oxt or PFC (downregulated Gas5; upregulated Cyr61, Gm12840, Dcn, Man1c1, Sox2ot) but remained unaffected in females (data not shown). 3 genes (Apold btg2, ddx39b) were upregulated in both Hypo and PFC but unaltered in females (data not shown). We, therefore, focused on Ttr and carried out detailed functional analysis pertaining to TH signaling in our experimental regime.

### PPS incites persistent changes in Ttr gene expression in both sexes but in opposite manner

RT-PCR validation of the transcriptome data revealed unique brain region and sex biased diametrically opposite expression pattern of the only gene Transthyretin (Ttr) in adult mice cohort. Ttr was also amongst the topmost DEGs based on fold change and p value (Fig.1D volcano plot). In PPS induced SEagg males, Ttr mRNA showed a decrease of 0.23-fold in Hypo and a robust increase of 8-fold in PFC relative to control (Ctrl) males (Fig. 3B). On the contrary, adult females that did not show aggressive phenotype (SNAgg), Ttr mRNA expression pattern was opposite to males, being increased in Hypo (13.6-fold) and drastically reduced (0.55-fold) in PFC relative to control counterparts (Fig. 3C)

In order to understand whether this gene expression changes was persistent from peripubertal age, we analyzed Ttr mRNA in brain regions post 24h after PPS exposure. The direction of Ttr mRNA changes was similar at peripuberty in both the brain regions and sexes although there were minor differences in the extent. PPS caused drastic reduction in Hypo (0.40-fold) and increase in PFC of Ttr mRNA expression (22.3-fold) of males (Fig. 3D). In females, the changes were reverse being upregulated (13.12-fold) in Hypo and reduced (0.51-fold) in PFC of PPS mice relative to unstressed (NS) controls (Fig. 3E)

### TTR protein alters in spatial and cell type specific manner

Immunoblot analysis of TTR protein levels corresponded to its transcript pattern in both the sexes. TTR protein was reduced to 0.37-fold in Hypo and upregulated by 1.36-fold in PFC in PPS induced SEagg males, relative to control (Ctrl) animals (Fig. 3F). Uncropped immunoblots of TTR has been included (Fig 3-source data 1). Until now we were considering the changes in bulk tissue, therefore, we performed immunofluorescence to elucidate spatial and cell type specificity if any. In SEagg males, TTR protein intensity was significantly reduced in Hypo and signals were prominent in rounded donut shaped cells although the cell type remains unidentified (Fig. 3G). Of note, TTR protein expression was reduced throughout Hypo including dorsomedial, ventromedial and arcuate nucleus. Therefore, in later stereotaxy experiments, coordinates were chosen (Materials and Methods) to cover all the above mentioned areas. In PFC, TTR protein intensity was markedly increased in the endothelial cells. (Fig. 3H). TTR was found unaffected in choroid plexus region, referred to as the main site of the protein synthesis (Supplementary Fig. S3). Of note, TTR showed changes in cells involved in uptake of blood borne substances in the brain which further intrigued us to check the TTR mediated uptake of thyroxine in the brain regions.

### TTR perturbation affects long term availability of thyroid hormone in brain with concomitant changes in transporters, deiodinases and target gene expression

To explore the functional consequences of perturbed TTR expression, we measured peripheral as well as brain region specific T4 and T3 content in both sexes. Circulating TH including total T4 and T3 in serum (Fig 4B-E) was neither altered in adulthood nor at peripubertal age in both sexes. Interestingly, brain TH content was remarkably altered corresponding to TTR gene expression right from peripuberty till adulthood. In adult SEagg males, total T4 and T3 was reduced in Hypo but increased in PFC as compared to control samples (Ctrl) (Fig. 4F, 4G). These changes of hypothalamic and PFC T4 and T3 content was persistent from early peripubertal (NS vs PPS males) age (Fig. 4H, 4I).

Besides TTR, local TH availability in the brain is dependent upon TH transporters and deiodinase enzymes that determine the intracellular conversion of T4 to T3. Further, TH mediates its action by regulating expression of target genes. Therefore, we addressed possible changes in brain local TH signalling by analysing levels of TH transporters, Mct8 and Oatp1c1 and deiodinase (Dio2). We also explored TH responsive genes that was differentially expressed in our transcriptome data (Trh, Nrgn). Both Mct8 and Oatp1c1 transcript levels were reduced in Hypo of SEagg males to 0.62-fold and 0.70-fold respectively, but remained unaltered in PFC as compared to Ctrl males (Fig. 4L, 4M). On the other hand, Dio2 mRNA showed an increase of 2.16-fold in Hypo but did not show significant change in PFC (Fig. 4K). DIO2 enzyme concentration was also increased in Hypo (Ctrl-11.1 ng/ml; SEagg-14.73 ng/ml) but was reduced in PFC of SEagg males (3.96 ng/ml) as compared to Ctrl (11.55 ng/ml) males (Fig.4J). In females, direction of changes for both T4 and T3 levels were reverse being increased in Hypo and reduced in PFC (Supplementary Fig. S4) but remained unaffected in serum.

Hypothalamic reduction in T4 and T3 content and consequent impaired TH signaling was clearly evident from expression of downstream target genes. TH responsive Nrgn mRNA expression showed significant downregulation of 0.5-fold in SEagg males compared to Ctrl males (Fig. 4O) in Hypo similar to Ttr mRNA. NRGN protein level was also reduced to 0.6-fold in hypothalamus of Eagg males (Fig.4P). Uncropped images of NRGN western blot has been included (Fig. 4-source data 2). Another, TH regulated gene, Trh showed a robust increase of 5-fold in Hypo of SEagg males while remained unaltered in PFC (Fig. 4N). Both Nrgn and Trh mRNA levels showed similar expression profile in early life peripubertal age (Supplementary Fig. S4) indicating a long term change in gene expression.

### Hypothalamus targeted TTR gene ablation impaired TH signaling and induced escalated aggressive behavior in males without peripubertal stress exposure

We checked the direct causal role of TTR by blocking its gene expression through jet-PEI mediated Ttr esiRNA injection in hypothalamus. Hypothalamus targeted Ttr knockdown to 0.2-fold (80% reduction) in adult unstressed males mirrored the escalated aggression induced by PPS (Movie V1-Scrambled injected mouse and Movie V2 Ttr siRNA injected mouse). Hypothalamus specific Ttr deficient males showed very short average attack latency of ~15 seconds and spent 60.5% of total behavioral RI session in clinch attack while none of the scrambled control animals showed signs of attack (Fig.5B and source file Fig 5). Such behavioral profile mirrored the phenotype of PPS exposed SEagg male cohort as shown in Fig1.

**Fig.5.**
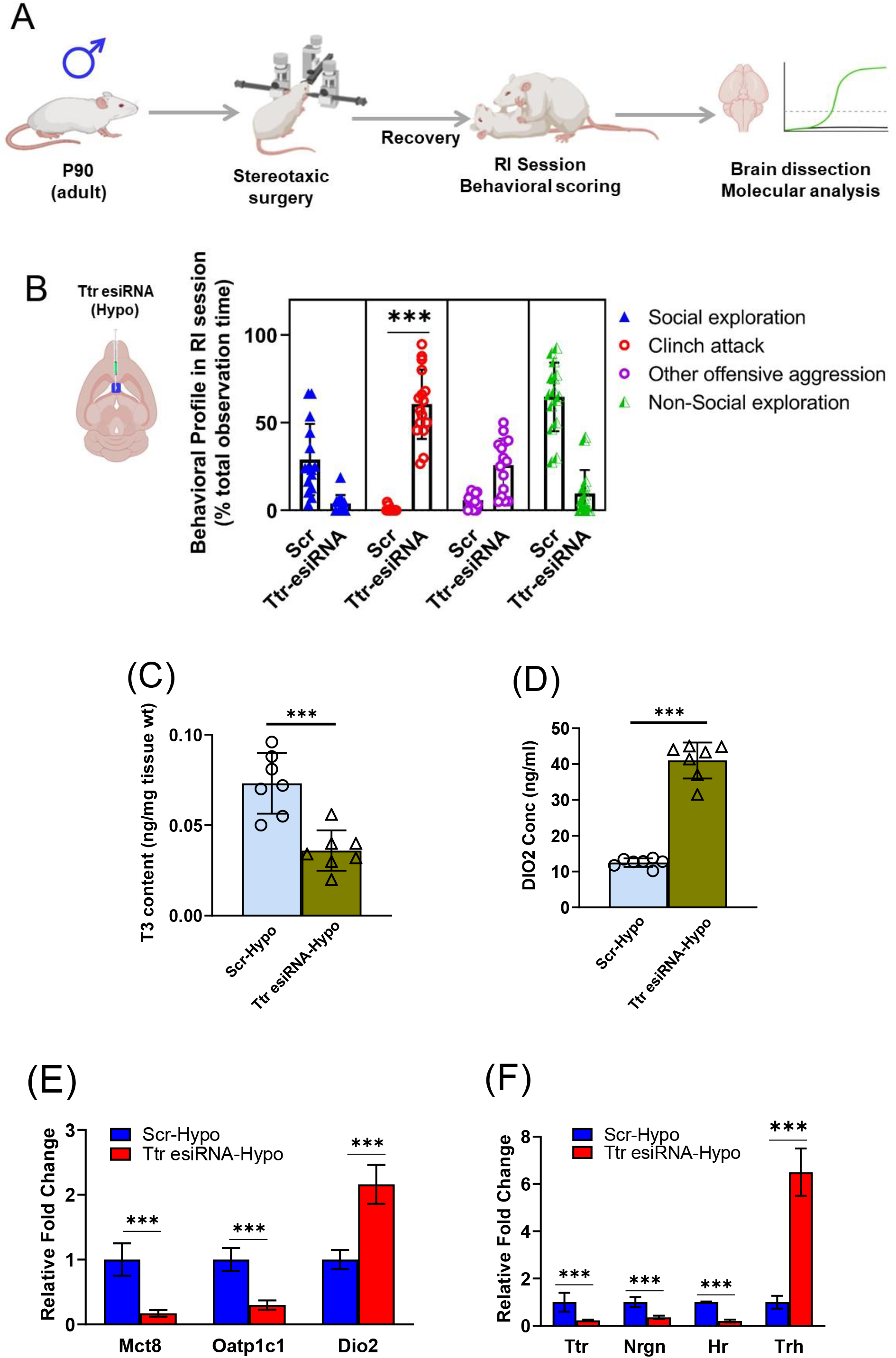
(A) (i) Experimental strategy of stereotaxic surgeries followed by behavioral and molecular experiments (ii) Diagrammatic representation of injection site in hypothalamus for JetPEI mediated Ttr esiRNA and scrambled siRNA (Scr) delivery. (B) Comparative analysis of behavioural profile during RI session between Ttr esiRNA (N=16 mice per group from 3 independent experiments and cohorts of mice) and Scr (N= 15 mice per group from 3 independent experiments and cohorts of mice) administered males. (C) T3 content in Hypo of Ttr esiRNA (N=7 mice per group from 2 independent experiments and cohorts of mice) and Scr males (N=6 mice per group from 2 independent experiments and cohorts of mice). (D) DIO2 enzyme concentration in Hypo of Ttr esiRNA (N=7 mice per group from 2 independent experiments and cohorts of mice) and Scr males (N=6 mice per group from 2 independent experiments and cohorts of mice). (E) Mct8 mRNA, Oatp1c1 mRNA and Dio2 mRNA expression and (F) Ttr mRNA, Nrgn mRNA, Hr mRNA and Trh mRNA expression analysis in Hypo of Ttr esiRNA and Scr males (N=9 mice per group from 3 independent experiments and cohorts of mice) Histogram represents mean of the data (±SD) Statistical analysis were performed using unpaired Student’s t-test [*** (p< 0.001)] between Scr vs Ttr esiRNA group.

T3 content in hypothalamus was decreased from 0.07 ng/mg tissue wt in scramble treated group to 0.03 ng/mg tissue wt in Ttr esiRNA treated group (Fig.5C and source file Fig 5). DIO2 enzyme levels showed pronounced increase from 12.5 ng/ml in scramble treated group to 41 ng/ml in Ttr esiRNA treated group (Fig.5D and source file Fig 5).Ttr gene silencing dramatically reduced mRNA expression of other transporters Mct8 and Oatp1c1 to 0.17 and 0.30-fold respectively. Dio2 mRNA showed an increase of 2-fold upon Ttr gene deficiency in hypothalamus (Fig.5E). TH regulated Trh mRNA was also markedly increased by 6.5-fold and Nrgn mRNA got reduced to 0.36-fold upon Ttr gene silencing. Here we included another well-established TH responsive gene hairless (Hr) that showed maximal downregulation to 0.2-fold upon Ttr gene silencing in hypothalamus (Fig.5F). Chemically induced increase in T4 levels by injecting levothyroxine in PFC did not show any significant behavioral changes (Supplementary Fig S5).

### Escalated aggressive behavioral phenotype is inherited in F1 males with impairment in Ttr gene expression and TH signaling in hypothalamus

We investigated whether PPS-triggered aggression of mouse strain Balb/c is propagated in next generation. Adult Eagg males that were exposed to PPS paradigm were mated with non-stressed females to generate the F1 progenies. F1 male and female progenies were examined at their adulthood. F1 male progenies of F0 Eagg males showed similar behavioral phenotype that characterized the parental generation including short attack latency, attack towards anesthetized and female intruder. F1 male progenies of F0 Eagg males spent 45% of RI observation time in clinch attack with extremely short attack latency of 11 seconds while males from control F0 did not exhibit attack (Fig.6B, 6C and source file Fig 6).However, female siblings of F1 males did not display any prominent sign of aggression.

**Fig.6.**
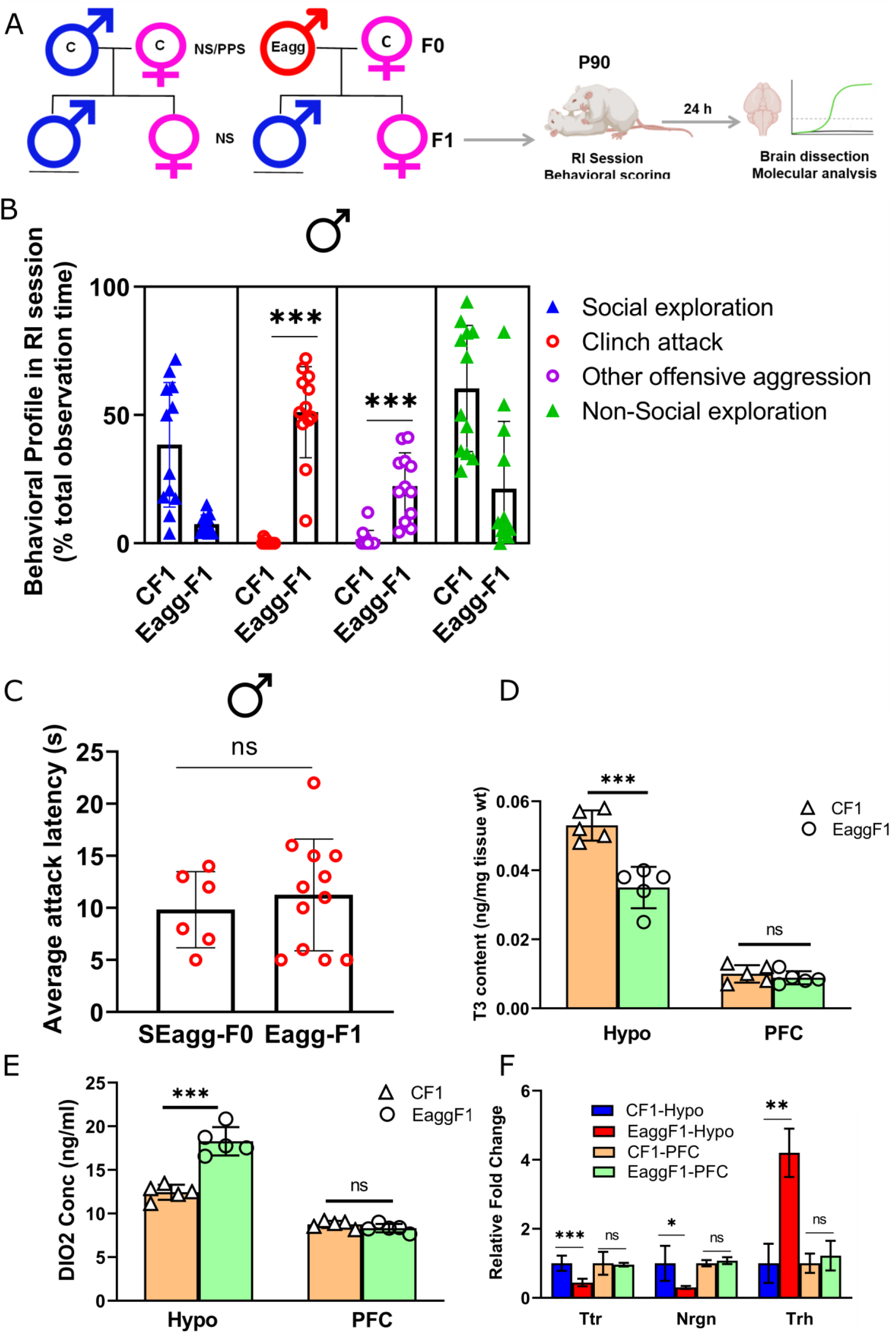
Intergenerational inheritance of PPS induced escalated aggression with concomitant changes in Ttr and thyroid hormone signaling. (A) Breeding pairs and experimental timeline. (B) Comparative analysis of behavioral profile during RI session between F1 males originating from control males crossed with control females (CF1) and F1 males originating from peripubertal stress exposed SEaggF0 males crossed with control females (EaggF1) males (N=12 mice per group from 2 independent experiments and cohorts of mice). (C) Attack latency comparison between parent SEagg male (SEaggF0; N=6 mice) and Eagg F1 male (EaggF1; N=12 mice). (D) T3 content in Hypo and PFC of CF1 and EaggF1 males (N=5 mice per group from 2 independent experiments and cohorts of mice). (E) DIO2 enzyme concentration in Hypo and PFC of CF1 and EaggF1 males (N=5 mice per group from 2 independent experiments and cohorts of mice). (F) Ttr, Nrgn and Trh mRNA expression analysis in Hypo and PFC of CF1 and EaggF1 males (N=6 mice per group from 2 independent experiments and cohorts of mice). Histogram represents mean of the data (±SD). Statistical analysis were performed using unpaired Student’s t-test [ns (p> 0.05),* (p< 0.05) and ** (p< 0.01)] between CF1 and EaggF1 groups or EaggF0 vs EaggF1.

Next, we checked whether molecular changes in parental generation (F0 Eagg father) including impaired Ttr gene expression and local TH signaling in brain were also perpetuated in the next generation. Similar to F0 Eagg father, F1 males showed deficiency in hypothalamic T3 content while that of PFC was not altered (Fig.6D and source file Fig 6). DIO2 enzyme levels also increased in hypothalamus but remained unchanged in PFC (Fig.6E and source file Fig 6). Ttr expression reduced to 0.35-fold in the hypothalamus of F1 Eagg males without any significant change in the PFC. Nrgn (reduction to 0.35-fold) and Trh (upregulation by 4.3-fold) were also altered similarly in hypothalamus (Fig.6F)

### PPS elicits long lasting change in Ttr DNA methylation in the brain of males showing escalated aggression

Next, we examined whether epigenetic regulation of Ttr could explain the sustained molecular and behavioral changes invoked by PPS exposure. To address this question, we analyzed DNA methylation state of Ttr proximal promoter in the PFC and Hypo of male mice. As anticipated, MedIP PCR demonstrated that peripubertal traumatic experiences trigger changes in Ttr DNA methylation within the PFC and Hypo in adulthood and even inherited in next generation. Ttr promoter showed brain region specific differential methylation state in opposite direction to that of its expression pattern. Hypermethylation was induced in Hypo (9.79-fold) where as hypomethylation (0.28-fold) was evident in PFC of PPS exposed escalated aggressive adult male of F0 generation (SEaggF0) relative to control (CF0). Hypermethylation in Hypo even persisted in the brain of F1 males that showed escalated aggression without PPS exposure (EaggF1) (5.7-fold) as compared to control males of F1 generation (CF1). In an independent study on DNA methylome of SEaggF0 male cohort (data not shown), we found Ttr as one of the topmost differentially methylated gene. As animals used for behavioral experiments were same as those for epigenetic studies, we inferred that Ttr promoter methylation could serve as a predictor of early life trauma induced gene expression and behavioral deficits (Fig. 7 and source file Fig 7)

**Fig. 7.**
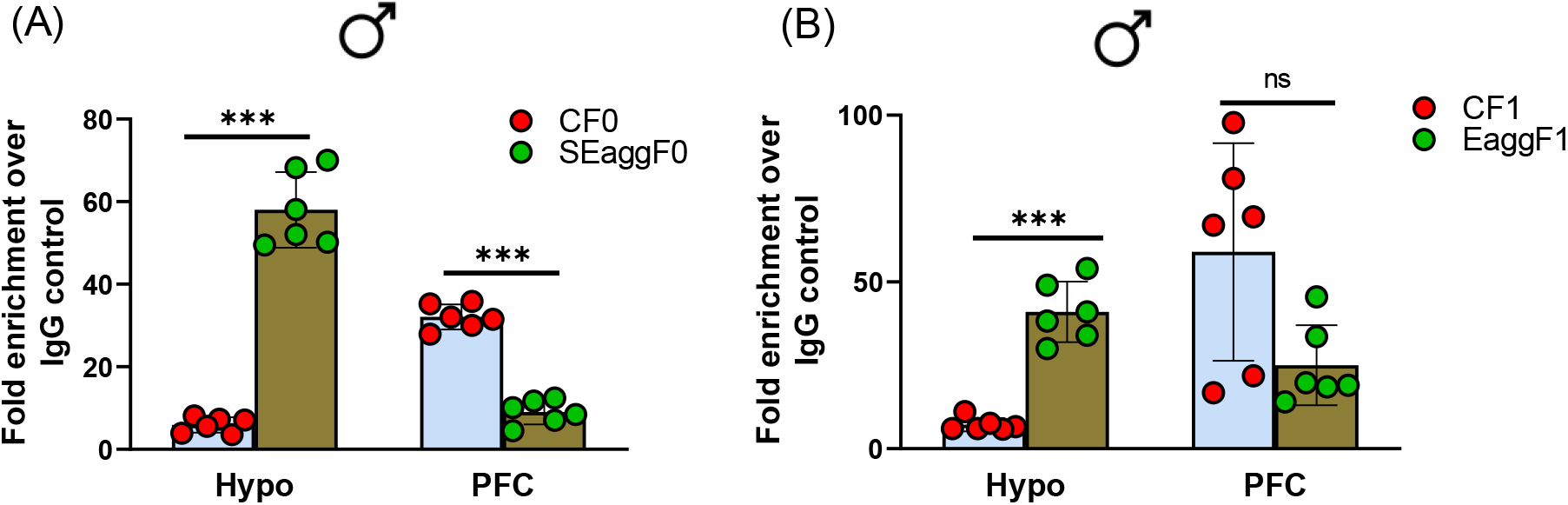
Ttr promoter methylation changes in F0 violent males perpetuated in F1 generation. Methylated DNA immunoprecipitation analysis showing 5-methylcytosine fold enrichment in Ttr promoter in Hypo and PFC of (A) Control male mice of F0 generation (CF0) and PPS induced male mice showing escalated aggression of F0 generation (SEaggF0) (N=6 mice per group from 2 independent experiments and cohorts of mice). (B) Control male mice of F1 generation (CF1) and F1 male mice originating from SEaggF0 and showing escalated aggression (EaggF1) (N=6 mice per group from 2 independent experiments and cohorts of mice). Histogram represents mean of the data (±SD). Statistical analysis were performed using unpaired Student’s t-test [ns (p> 0.05) and *** (p< 0.001)] between CF0 and SEagg F0 or CF1and Eagg F1 males.

## Discussion

In the present study, we fill in the existent gap of knowledge about the molecular roots of escalated aggressive behavior. Based on unbiased transcriptome screening, we identify novel role of TTR in long-term programming of escalated aggression induced by PPS. More importantly, we show that TTR regulated thyroid hormone availability in hypothalamus contributes to abnormal behavioral response.

TTR is a 55 kDa protein that is involved in the uptake of T4 from blood to CSF and local distribution of the hormone in brain (*13*). TTR has also been assigned other functions including proteolysis of Neuropeptide Y (*14*), neuroprotection and regeneration of damaged neurons (*15*). However, we focused on TH based on multiple reasons. Our transcriptome data showed significant expression change in TH genes primarily Trh and Nrgn in hypothalamus. Several other genes in our list (PFC-Rasd2, Fosl2, Nr4a3, Inf2, Arl4d, Dcn, Pcp4l1, Drd2, Syndig1l, Hspa1a, Spock3; Hypothalamus-Oxt, Cdhr1, Col23a1, Dgkk, Dkk3, Cck, Ptpro;) showed overlap with literature available on T3 responsive genes in primary cultured neurons (*16*, *17*). Moreover, we observed persistent TTR expression change right from peripubertal age and perturbation of TH action in developing brain rather than adult stage have been considered as critical determinants of multiple neurological deficits (*18*). TTR known function in TH transport and distribution as well our finding of its alteration in brain endothelial cells involved in uptake of blood borne substances into brain (*19*, *20*) indicated that transport functions might be hindered in our experimental paradigm. It is important mentioning here that we showed for the first time that TTR is expressed in PFC endothelial cells and hypothalamus donut shaped cells, cell type remaining to be identified. Earlier studies report TTR synthesis in choroid plexus epithelial cells of the brain until recently it was identified in neurons (*21*, *22*) and astrocytes indicating wide expression of the protein in CNS.

In clinical settings, thyroid hormone abnormalities are diagnosed by serum parameters. However, we show that alterations in TTR gene expression paralleled perturbation in T4 and T3 content in brain tissues of hypothalamus and PFC without affecting the circulating levels of the hormone. Determining the effective concentration of T4 and T3 in brain tissues is difficult owing to multiple factors driving their synthesis, transport across blood-brain and blood-CSF barrier, intracellular distribution and activation/inactivation (23, 24). Therefore brain specific alterations in TH intrigued us to explore factors besides TTR including transporters Mct8, Oatp1c1 and enzyme Dio2 that determine the local TH availability in brain (25, 26).

We noted significant downregulation of Mct8 and Oatp1c1 mRNA in hypothalamus of PPS induced escalated aggression showing males that corresponded well with reduced hypothalamic T4 and T3 content. Previously, it was reported that mutation in thyroid transporter MCT8 gene or decrease in its expression hindered intracellular T3 distribution and neuro-glia communication eventually leading to neurological ailments like multiple sclerosis (*27*). Also, Mct8 knockout mice showed pronounced deficiency in brain T3 content and was region specific particularly hypothalamus but not in other areas (28)

Mct8 deficiency in rodents is considered to be compensated by T4 transport through Oatp1c1. Oatp1c1 was also reduced in our model system suggesting significant local hypothyroidism in hypothalamus owing to lack of both the transporters (25, 29). In particular, tissues such as brain that depend only on these transporters for TH uptake and distribution, become hypothyroid upon their deficiency whereas other tissues including liver and kidney can still maintain levels of T3 (30).

Dio2 levels showed subtle increase in hypothalamus of PPS induced violent males. In brain, T3 is derived partly from circulation and is also formed locally by Dio2 mediated 5’-deiodination of T4. Dio2 action is integral to ascertain optimal intracellular concentrations of T3 (31). Therefore, Dio2 increase is suggested to be a compensatory response to deficiency of both Mct8 and Oatp1c1 and maintain TH levels (32). Dio2 is known to be negatively regulated by TH and was increased upon hypothyroidism in aging rodent brain (33). This literature information indicated that DIO2 increase in hypothalamus in our model system strongly suggests reduction in local TH availability.

Any change in local TH status of brain has a direct influence on expression of TH responsive genes. T4 and T3 action initiates with formation of ligand-receptor complexes with TH nuclear receptors (TRs) which in turn binds to TH response element (TRE) on the promoter of target genes. A broad spectrum of TH responsive genes are critical for plethora of TH action in varied cellular processes such as cell proliferation, differentiation, metabolism and homeostasis (34,35). TH deficiency during postnatal brain development causes irreversible neurological manifestations through target gene expression changes (36).

Nrgn is one such brain specific TH responsive gene that was also amongst the top ranking differentially expressed gene in our transcriptome data containing TRE elements in promoter and its transcription is dependent on TH in brain (37,38). Nrgn regulates synaptic plasticity by activating calmodulin kinase II (CaMKII) protein and spine density. We observed significant reduction in Nrgn transcript and protein levels in concordance with hypothalamic decrease in T4 and T3 levels. Another top ranking gene of our transcriptome data, Trh known to be negatively regulated by T3 was robustly upregulated in hypothalamus of males showing escalated aggression. Overall these findings implicated the role of reduced local thyroid hormone availability in hypothalamus in PPS induced escalated aggression.

Until now we found a strong link between reduced local TH availability in hypothalamus and emergence of escalated aggressive phenotype but whether it was mediated by Ttr or was independently regulated was not clear. We showed that intra-hypothalamic Ttr gene knockdown led to similar reduction in T3 availability, decrease in Mct8 and Oatp1c1, increase in Dio2 and alteration in expression of TH target genes as found in our earlier experiments.

Besides Trh and Nrgn, Ttr silencing also reduced hairless (Hr), a universal TH responsive gene that is studied to monitor the local TH status in brain (39). Therefore, we included Hr mRNA expression analysis in our study. Ttr gene ablation also evoked escalated aggression in unstressed males to a similar extent to that of PPS induced males. These data clearly indicated that behavioral and molecular consequences in our experimental regime was downstream of Ttr, though detailed studies are required to delineate the precise mechanism of action of Ttr in hypothalamic cells.

Interestingly, TTR showed expression changes in both hypothalamus and PFC whereas escalated aggressive behavior was triggered only by hypothalamic TTR knockdown. TTR overexpression or T4 increase by levothyroxine in PFC did not produce any behavioral changes (data not shown). It is likely, that TTR mediated thyroxine reduction in hypothalamus have direct causal role in aggression and consequently PFC endothelial cells express more TTR to uptake hormone and alleviate the deficiencies. Interestingly, we did not observe significant change in other transporters Mct8 and Oatp1c1 in PFC further indicating that TTR mediated TH uptake in PFC might result in the compensatory increase of T4 and T3 in response to hypothalamic reduced TH availability.

We explored intergenerational inheritance of PPS induced escalated aggression and observed paternal transmission of behavior in F1 males with concomitant reduction in, hypothalamic TTR and Nrgn expression and T3 availability. Previous studies suggest that thyroid hormone changes in neonatal brain can elicit neuroendocrine abnormalities in their F1 progenies. Also, developmental exposure of thyroxine disrupting chemicals can affect gene expression and behavior in later generations (40). Further, mechanistic investigation revealed lasting methylation mark in TTR promoter which was inherited in F1 generation. DNA methylation plays crucial role in the inheritance of traumatic memories (41, 42). Therefore, we speculate that epigenetic inheritance at the TTR methylation locus controls long-term programming of the hypothalamic-thyroid axis which in turn modulates thyroid hormone availability and function throughout life and in subsequent generations. Future epigenetic manipulation at TTR locus during PPS can shed light on life-long vulnerability to pathological aggression and its inheritance.

In conclusion, we provide molecular evidence for emergence, gender vulnerability and inheritance of post traumatic escalated aggressive behavior. We delineate novel role of brain TTR-thyroid signaling in manifestation of early life trauma induced excessive aggression and its intergenerational inheritance that could be mediated at least in part by epigenetic mechanisms. Brain TTR-thyroid signaling can also serve as valid molecular predictors as well as intervention targets in excessive pathological aggression. Our findings have inherent limitations of investigations in animal models and therefore further studies are warranted in relevant human cohort to establish role of TTR-thyroid signaling pathway in abnormal aggression and related psychopathologies. Our work also provides resource for investigating sexual dimorphism in behavioral disorders and deciphering susceptibility as well as protective pathways.

## Materials and Methods

### Animals

All experimental procedures involving live animals were approved by the Institutional Animal Ethics committee (IAEC) of CSIR-Institute of Genomics and Integrative Biology (IAEC Approval Number-IGIB/IAEC/3/15) that is registered under Committee for the Purpose of Control and Supervision of Experiments on Animals (CPCSEA), Department of Animal Husbandry and Dairying, Ministry of Fisheries, Animal Husbandry and Dairying, Government of India (Registration No and Date-9/1999/CPCSEA). Male and female offspring of Balb/c mice bred in the institutional animal house were used for the study. All animals were housed under SPF conditions. They were kept in individually ventilated cages (IVC) at 24±2°C on a 12h light/dark cycle with ad libitum access to food and water. Animal handling and experiments were conducted in accordance with the institutional guidelines.

### PPS stress procedure

Male and female mice were exposed to unpredictable fear inducing stressors of synthetic fox odor (trimethylthiazoline) and elevated platform during the peripuberty period of postnatal day (P) 28 to P42 as per the protocol published previously (6,7). Post weaning at P21, equivalent number of mice from different litters were mixed and placed in control and experimental (peripubertal stress-PPS) groups in different home cages (3-4 mice per cage) avoiding placing of siblings in the same home cage.

Briefly, P28 male and female offspring were exposed to an open-field for 10 minutes for acclimatization in a novel environment. Thereafter, one group of mice were exposed to 9 μl of fox odor (Sigma) soaked cloth kept in a filter top plastic cage and elevated platform (96 cm above ground) for 7 random days (P28, P29, P30, P34, P36, P40 and P42) across P28 to P42. Stressors were applied during the active phase of the mice, singly or in combination in variable schedule so that the animals do not learn and get suddenly traumatized. The duration of stress session was 25 minutes following which mice were returned to their home cages. Control animals were handled on the days in which their counterparts were exposed to PPS.

### Resident intruder (RI) paradigm

Control and PPS exposed mice of both sexes were assessed for aggression in their adulthood (P90) using the conventional RI paradigm. Mice were individually housed for 1 week prior to testing and RI test was performed in active phase of the animals.The resident was exposed in its home cage to various category of intruders including a smaller and unfamiliar (10% less body weight) size, larger size (10% more body weight), anesthetized, opposite sex and intruder of different strain for 10 minutes for 7 consecutive days. Each day the resident was introduced to a different intruder in a latin square design. The behavioral parameters including clinch attack, move towards, social exploration, ano-genital sniffing, rearing, lateral threat, upright posture, keep down, chase, non-social explore and rest or inactivity were quantified in terms of percentage (duration) of the total observation time. Attack latency or the time between introduction of the intruder and first clinch attack was also determined. The total duration of the clinch attack, offensive upright, keeping down and lateral threat were considered as the measure of total offensive behavior. Social exploration behavior included the sum of social explore, auto and social grooming and ano-genital sniffing. Phenotypic screening of animals into non-aggressive, less-aggressive, moderate-aggressive and hyper-aggressive was done based on conventional parameters as published in earlier reports (12, 43). A sub cohort of hyper-aggressive animals showing extreme phenotype was referred to as escalated aggressive. The order of RI testing for control and PPS exposed adult animals was random. Behavioral scoring was done by an observer blind to animal identity and assignment of groups.

### Breeding scheme for F1offspring

Control (without PPS exposure) and PPS exposed adult male mice showing escalated aggression was mated with control females (without PPS exposure). After mating, males were immediately removed from the cage so that they do not have any contact with their offspring and do not impact upon their rearing. F1 offspring originating from these pairings were housed in standard cages and subjected to RI test for aggression at their adulthood (P90).

### RNA-sequencing

RNA was isolated from hypothalamus and PFC of male and female mice using Trizol reagent. Approximately, 1μg of RNA was taken per sample and RNA sequencing libraries were made using TruSeq v2 Library Prep Kit as per manufacturer’s protocol. Briefly, the RNA was polyA selected using OligodT magnetic beads followed by shearing into 200-500 bp fragments. This sheared RNA was then used to generate cDNA. The cDNA was end-repaired to blunt ends. These blunt ends were then A-tailed i.e. an “A” overhang was added so as to ligate the adapters in the next step. The adapter-ligated cDNA was then amplified by PCR and purified by AMPure XP beads. The prepared library was quantified using Qubit Fluorometer, and validated for quality on High Sensitivity Bioanalyzer Chip (Agilent) and sequenced on Illumina HiSeq 2500. The FASTQ sequencing reads were adapter-trimmed along with a minimum length cut-off of 50 bases using Prinseq-lite. The reads were aligned to mouse genome assembly using TopHat (v.2.0.11) followed by reference-based assembly using Cufflinks (v.2.2.1). Then differentially expressing transcripts were identified using Cuffdiff (v.2.2.1). Raw and processed RNA-seq datasets were deposited in National Center for Biotechnology Information (NCBI) Gene Expression Omnibus (GEO), accession number GSE199844.

### RT-PCR

Total RNA was isolated from hypothalamus and PFC of mice and 2 μg of RNA from each group was reverse transcribed to cDNA synthesis. RT-PCR was carried out using SYBR Green master mix for detection in Light cycler LC 480 (Roche). All primers used for qRT-PCR are given in Supplementary Table S1.The endogenous control GAPDH was used to normalize quantification of the mRNA target.

### Immunoblotting

Cytosolic protein lysates (40 μg) prepared from mouse hypothalamus and PFC were resolved on to 10% SDS PAGE, transferred to PVDF membrane and used for immunoblotting using conventional method. The primary antibodies {anti-TTR rabbit polyclonal, anti-Nrgn rabbit polyclonal; anti-GAPDH mouse monoclonal} and secondary antibodies {anti-rabbit IgG HRP (Cell Signaling Technology, 7074P2) and anti-mouse IgG HRP (Cell Signaling Technology, 7076P2)} were used at adequate dilutions.

### Immunohistochemistry

Mice were anaesthetized with thiopentone (40 mg/kg) and perfused with cold 4% paraformaldehyde in PBS. Brains were removed, post-fixed, cryoprotected in PBS + 15% sucrose for 2–3 hours followed by immersion in PBS + 30% sucrose for 24 h, and then sectioned coronally (7 μm) on a cryotome. Free-floating sections were permeabilized with blocking buffer (PBS + 3% normal donkey serum, 0.3% Triton X-100) for 2 hours and then incubated with TTR primary antibody overnight at 4°C. Slices were then washed 4×15 min with PBS, incubated with corresponding secondary antibodies for 2 hours, washed 4 × 15 min with PBS, mounted on microscope slides followed by counterstaining with DAPI and photomicrographs were captured by FLoid fluorescence microscope.

### Thyroid hormone measurement

Mouse blood samples were collected from heart to test serum levels of total tetraiodothyroxine (T4) and total tri-iodothyroxine (T3). Thyroid hormone content in brain regions was determined by dissecting hypothalamus and PFC and individually homogenizing them in artificial cerebral spinal fluid and centrifuged at 14,000 rpm for 15 min at 4°C. The resulting supernatant was collected and used for ELISA based determination of total T4 and T3 (EliKineTM Thyroxine (T4) ELISA Kit KET007 and EliKineTM Triiodothyronine (T3) ELISA Kit KET006).

### Deiodinase 2 (DIO2) measurement

We performed an in vitro quantitative measurement of DIO2 in mouse brain tissue (hypothalamus and PFC) homogenates using a sandwich enzyme immunoassay kit (Reddot Biotech INC., Mouse Deiodinase, Iodothyronine, Type II (DIO2) ELISA Kit RDR-DIO2-Mu). Briefly, mouse hypothalamus and PFC tissues were isolated from control and experimental groups, homogenized in 1XPBS. The resulting suspension was subjected to 2 freeze/thaw cycles to break the cell membranes and centrifuged for 5 minutes at 5000 × g. The supernatant was removed and used for DIO2 ELISA assay as per manufacturer’s instructions. The concentration of DIO2 was measured spectrophotometrically using a microplate reader at a wavelength of 450 nm.

### Stereotaxic surgeries and gene manipulation

Mice were anesthetized with 40 mg/kg BW thiopentone i.p. and positioned on a robotic stereotaxic frame (Cat no. 51700, Stoelting Co., USA) with motorized stereo-drive (Cat no. 013.641, Neurostar, USA). As mentioned in earlier reports (44), the dorsomedial, ventromedial and arcuate nucleus of hypothalamus were targeted bilaterally by using the stereotaxic coordinates of 1.5 mm posterior to the bregma, 0.5 mm lateral to midline, and 5.8 mm below the surface of the skull. For PFC, the specific coordinates for injection relative to bregma was mediolateral ± 0.35mm, dorsoventral −2.1mm, and rostrocaudal axes +2.2 mm. For brain targeted gene manipulation TTR esiRNA (esiRNA targeting mouse Ttr- EMU030721, Sigma Aldrich)–jetPEI complex was infused into hypothalamus. Levothyroxine (LT4) was bilaterally administered into PFC. Injection was given using 10μl Hamilton syringe (Cat no. 72-1823,32G; 700N glass) placed in arm-held Elite-11 mini pump (Harvard Apparatus, USA) at a rate of 100 nl/min and the system was left in place for an additional 1 min and then gently withdrawn. Mice were allowed to recover individually from anesthesia and thereafter returned to their home cages. Post-operative cares were taken using analgesics and anti-biotics including meloxicam (5mg/kg of b/w, i.m., Intas Pharmaceuticals, India) and gentamycin (5mg/kg bw, i.m., Neon Laboratories, India) for 2-3 days. Body temperature was maintained during and after surgery in homoeothermic monitoring system (Harvard Apparatus, USA) following previous protocol. RI test for aggression was performed followed by molecular experiments after an additional 24 h.

### Methylated DNA immunoprecipitation

DNA methylation was analyzed at the promoter region of Ttr by methylated DNA immunoprecipitation (MeDIP) method as mentioned earlier (45, 46). Briefly, 4 μg of sonicated DNA (DNA fragment size ranging from 300 to 1000 bp) isolated from hypothalamus and PFC of F0 and F1 male mice was diluted in immunoprecipitation buffer and incubated with 2 μg 5-methyl cytosine antibody (A-1014; Epigentek) at 4°C overnight. Mouse IgG Isotype control antibody (02-6502, Thermo Fisher Scientific) was used for mock IP. Next day, 50 μL of Protein A-dynabeads was added and incubated at 4°C for 2h with rotation. Thereafter, it was centrifuged at 3500xg at 4°C for 10 min and the supernatant was removed carefully. After washing the pellet, the immune complex was eluted, DNA was purified and dissolved in TE buffer. Using eluted DNA as template, Ttr proximal promoter −184 to −33 bp from TSS) was amplified with specific primers (Supplementary Table 1) generating a 151 bp product in MedIP-qPCR.

### Statistical analyses

All statistical analyses were performed using Microsoft Excel or Prism 8 (GraphPad Software). Sample sizes were not predetermined during study design. However, for each experiment our sample sizes were based upon conventions established in previous publications including our own, that had used similar assays and power calculations. Sample sizes, statistical analysis used and exact p values for each experiment is mentioned in respective results and figure legend section and also uploaded as source files wherever relevant. Information regarding number of experiments and number of replicates are provided in each result and figure legend sections.

Total number of animals used in experiment and number of animals in each experimental group is also mentioned. Each animal was considered as a biological replicate and all our replicates presented in figures were biological. Experiment repeated with same samples was considered as technical replicate. In case of RT-PCR, immunoblotting, immunofluorecence, ELISA and MedIP experiments each biological replicate was further run in 3 technical replicates. All experiments except RNA sequencing (Figure 1 and 2) were repeated at least 2-3 times at least with independent cohort of mice. Experiments were repeated by separate researchers/experimenters wherever possible, to ascertain reproducibility and independent verification of data.

In order to analyze RT-PCR data, the 2^-ΔΔCt value was used to calculate relative fold change in mRNA expression and plotted as histograms. For immunoblot analysis, the signal intensity (Integrated Density Value, IDV) of TTR and NRGN bands was measured by spot densitometry tool of AlphaEaseFC software (Alpha Innotech Corp, San Jose, CA, USA), normalized against the IDV of internal control GAPDH and histogram was plotted as relative density value. For MeDIP analysis, results were represented as fold enrichment normalized to IgG control. Histograms were represented as mean of the data (±SD) and statistical significance was calculated by Student’s unpaired two tailed t-test.

## Acknowledgments

We acknowledge the animal house facility of CSIR-IGIB, New Delhi, India. We thank Ashish Kumar (Centre for Biomedical Engineering, IIT Delhi, India) for assistance in stereotaxy experiments.

## Funding

This work was supported by grants from Department of Science and Technology, Govt of India (DST/INSPIRE/04/2014/002261/GAP0125), Department of Biotechnology, Govt of India (GAP0197) and Indian Council of Medical Research (IR-594/2019/RS).

## Author contributions

A.K. conceived idea of the project with input from B.P. A.K. and R.R designed the experiments and interpreted the data. A.K., R.R., A.B., M.P., performed the behavioral, RNA sequencing and other molecular experiments. A.R. performed the stereotaxy surgeries with input from S.J. A.K. wrote the manuscript with input from B.P.

## Competing interests

The authors declare no competing interests, financial or otherwise.

**Movie V1**- Video showing resident intruder behavioural session in which resident mouse (legs marked) is scrambled siRNA injected in hypothalamus and intruder is a control mouse of lesser body weight. Scrambled injected mouse did not show any sign of offensive aggression and clinch attack.

**Movie V2**- Video showing resident intruder behavioural session in which resident mouse (body marked) is Ttr siRNA injected in hypothalamus and intruder is a control mouse of lesser body weight. Ttr siRNA injected mouse showed escalated aggression spending maximum time in clinch attack and biting during the entire behavioural session.

## Notes

### Competing Interest Statement

The authors have declared no competing interest.

