## Supplemental information for "Early life trauma leads to escalated aggressive behavior and its inheritance by impairing thyroid hormone availability in brain"

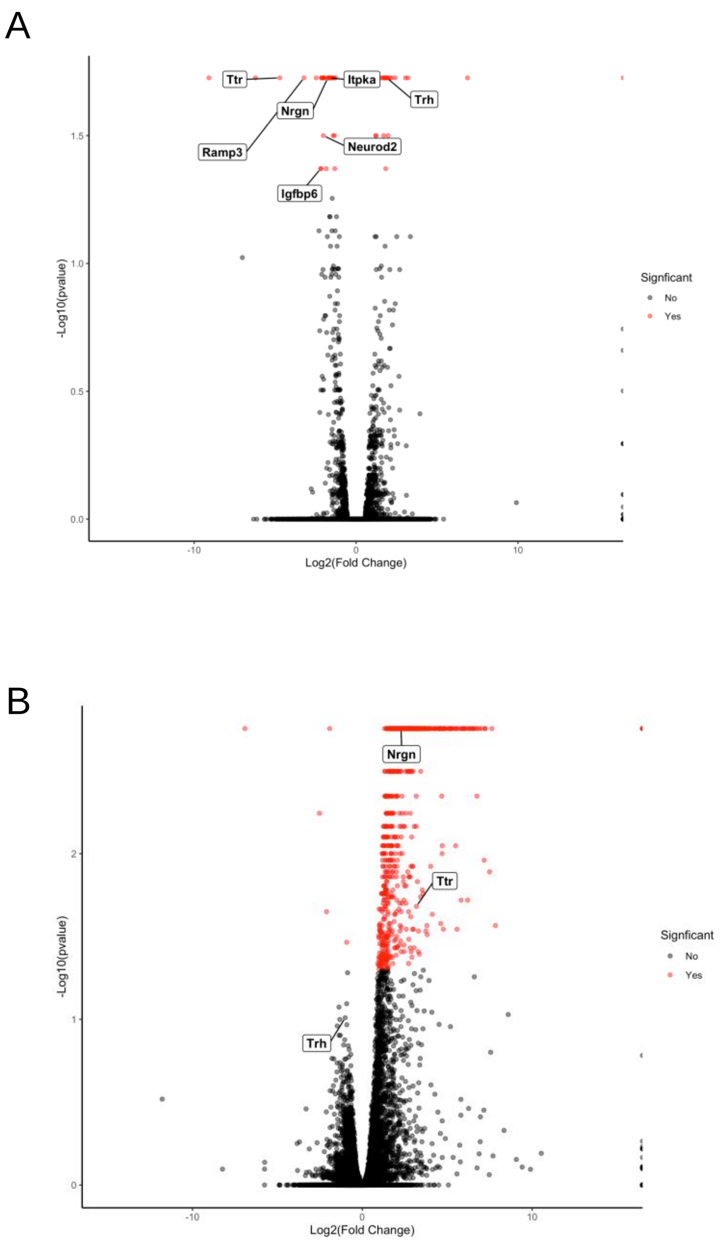


**Fig. S1 (**A) Volcano plot showing top ranking differentially expressed genes including Ttr, Nrgn and Trh in hypothalamus of PPS induced escalated aggressive males compared to control males. (B)Volcano plot showing Ttr, Nrgn differentially expressed in opposite pattern in PPS induced non-aggressive females compared to control counterparts

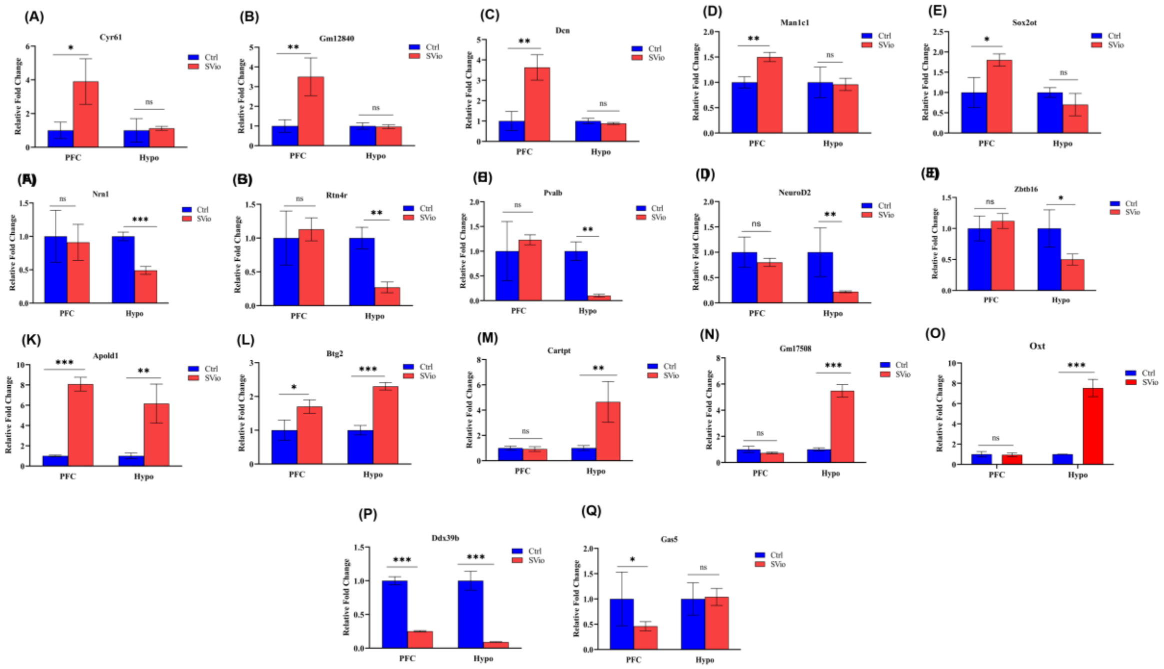


**Fig. S2.**RT-PCR validation of differentially expressed genes in PFC and hypothalamus of peripubertal stress induced escalated aggressive adult males. A-E, F-J , K-L and M-O genes show similar trend of expression pattern

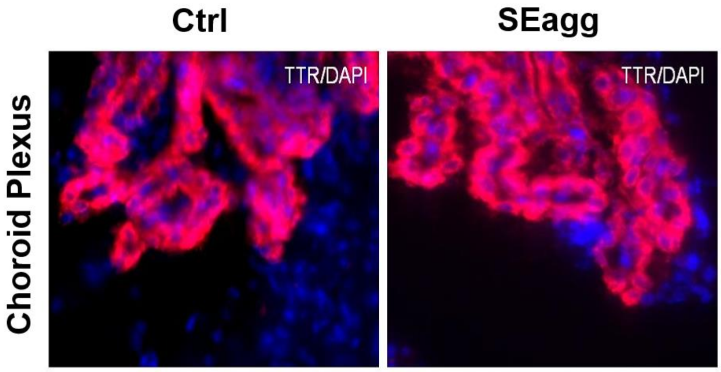


**Fig. S3.**TTR protein expression remained unaltered in choroid plexus of Ctrl and SEgg males.

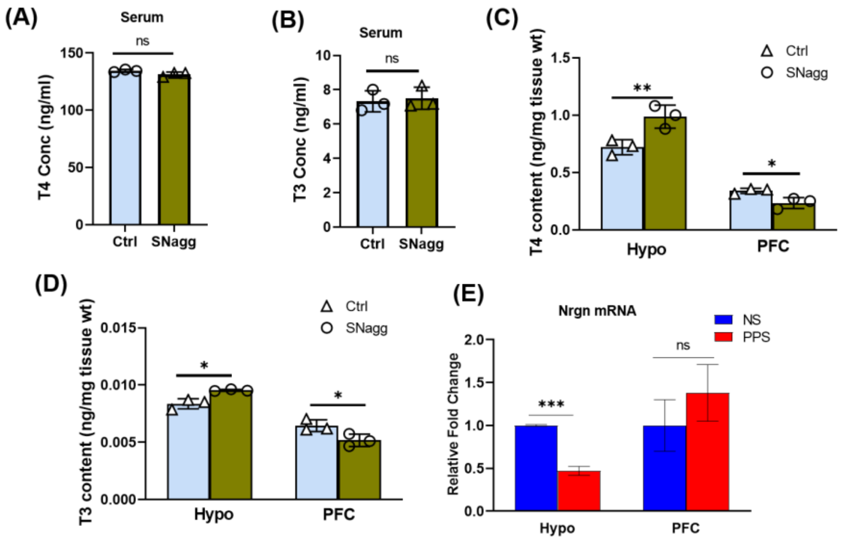


**Fig. S4**. (A) T4 and (B) T3 levels in serum and in brain regions PFC and Hypo (C,D) of Ctrl and SNagg females. (E) Nrgn mRNA expression in NS and PPS males.

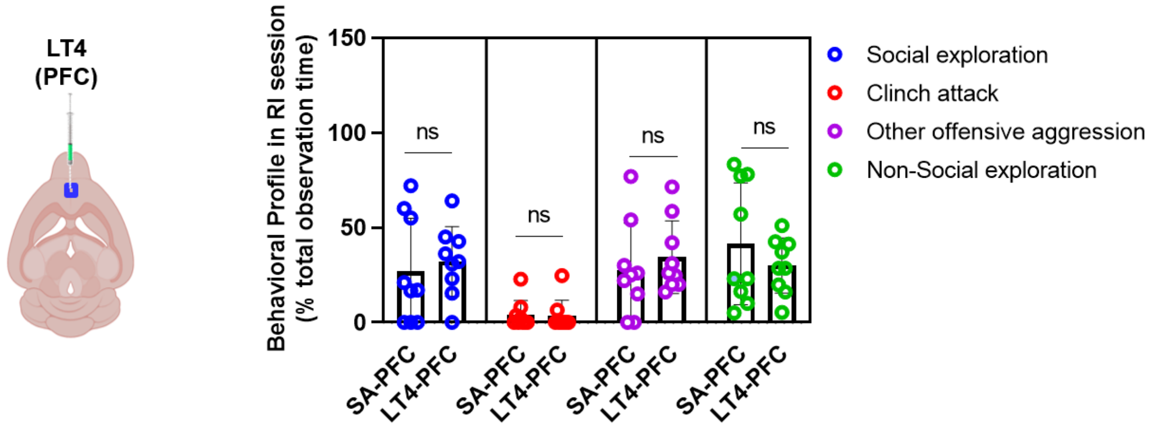


**Fig. S5**. Comparative analysis of behavioral profile during RI session between saline injected group in PFC (SA-PFC; N=9) and levothyroxine, LT4 injected in PFC (LT4-PFC;N=9). Histogram represents mean of the data (±SD) Statistical analysis were performed using unpaired Student's t-test [ns ($p > 0.05$)] between SA-PFC vs LT4-PFC group.

| **S.No** | **Gene** | **Forward Primer Sequence** | **Reverse Primer Sequence** |
|---|---|---|---|
| 1 | Apold1 | TGGAGGCCAGAGTGAAGG | CCAGCAGCAATCCTTGGAAG |
| 2 | Btg2 | GGGTTTCCTCTCCAGTCTCC | ACCTTGCTGATGATGGGGTC |
| 3 | Cyr61 | CCAATGACAACCCAGAGTGC | CCGCATCTTCACAGTTCTGG |
| 4 | Cartpt | GACATCTACTCTGCCGTGGA | CAGGACTTCTTGCAACGCTT |

| | | | |
|---|---|---|---|
| 5 | Dcn | ATCACCAAGCTGCGGAAATC | GGCTCCGTTTTCAATCCCAG |
| 6 | DDx39b | TCCAGCATGAGTGCATCCCG | AAAAGCCAGCTCCCTAGTGT |
| 7 | Dio2 | AGGATGCACACGGAAAGGAG | CCAGACCCCCTGGAGTTTTC |
| 8 | Gas5 | TTCATTTGGCTGGCTTGCTT | TTGTGTTTGCAGTGCCTTCA |
| 9 | Gapdh | TGTGTCCGTCGTGGATCTGA | CCTGCTTCACCACCTTCTTGA |
| 10 | Gm12840 | GTACAAGTCCAGGCGTGG | CAGCAGCCACAACATT |
| 11 | Gm17508 | TGAAGATATGGCACGCCCAG | AGCAAGGACACACACCAGTT |
| 12 | Hr | AAGCTAAATAGGGGATCCTG | ATTTGTAGAACGGACCACAC |
| 13 | Man1c1 | CCAACCATGACAACAGGCAA | GATGAGCCTCGGTGTTGAAC |
| 14 | Mct8 | GGGTAGGGGGCAGAATAAGTG | ACCCTGCCAATGTTAGGTGA |
| 15 | Nr4a | CTGTGGGCATGGTGAAGGAA | GAGGCTGCTTGGGTTTTGAA |
| 16 | Nrn1 | CACAGCTCTTACGGATTGCC | CAGAAGGAAAACCAGGTCGC |
| 17 | Nrgn | TAAAGAGCGGAGAGTGTGGC | AGGGGCTCACAAACACAGTA |
| 18 | Oatp1C1 | GCTGTGGAAAACTCAAGGTG | GGAAGGGATCTCAAACCTTC |
| 19 | Oxt | CCATCACCTACAGCGGATCT | CAGATGCTTGGTCCGAAGC |
| 20 | Pvalb | ATTGAGGAGGATGAGCTGGG | GTCCCCATCCTTGTCTCCAG |
| 21 | Rtn4r | TCCCTGACAACACCTTTCGA | ATGGTTCTGGTGCAAGAGGA |
| 22 | Sox2ot | ATCTCCAGGCAGAAGAGGAC | TCTGCTCAGTGGCTTCTCAA |
| 23 | Trh | TCGTGCTAACTGGTATCCCC | TCTCTTCGGCTTCAACGTCT |
| 24 | Ttr | GCTTCCCTTCGACTCTTCCT | TTACAGCCACGTCTACAGCA |
| 25 | Ttr Promoter | AGCGAGTGTTCCGATACTC | ACCCCCTCCTTCCAACCCA |
| 26 | Zbtb16 | AGTGCCTGAAGATCCTGGAG | TGGCTGAGAGACCGAAAGAG |

**Table S1: Primers used for qRT-PCR and qMeDIP.**
Gene name, forward and reverse primer sequence of primer pairs used in real-time PCR amplification of mRNA (qRT-PCR;) and 5 methyl cytosine immunoprecipitated DNA (qMeDIP)
